# Glutamatergic and Redox Abnormalities in Major Depressive Disorder: Links to Childhood Maltreatment and Perceived Stress

**DOI:** 10.64898/2026.10.28.762615

**Authors:** Sarah Syed, Rotem Dan, Lauren M. Hutson, Aliza R. Brown, Chao Wang, Abigail Stein, Xi Chen, Fei Du, Diego A. Pizzagalli

## Abstract

**Background:** Childhood sexual abuse (CSA) increases the risk for major depressive disorder (MDD), yet the underlying neurobiological mechanisms remain poorly understood. We examined (1) associations of medial prefrontal cortex (mPFC) glutamatergic metabolites and redox balance with childhood maltreatment and MDD; and (2) whether these neurochemical measures predicted future perceived stress.

**Methods:** Using proton and phosphorus 4T magnetic resonance spectroscopy, we quantified glutamate, glutamine, their sum (Glx), and the NAD /NADH ratio within the same mPFC voxel in 71 women across four groups: healthy controls (HC; *n*=27), current MDD without CSA (*n*=14), current MDD with CSA (*n*=14), and remitted MDD with CSA (*n*=16).

**Results:** Glutamate and Glx levels were lower in both CSA-exposed groups (*n*=30) compared with HC (*n*=27), whereas women with MDD without CSA did not differ significantly from HC. Among CSA- exposed women, greater maltreatment severity was associated with lower glutamine, and greater maltreatment multiplicity with lower glutamate and Glx, specifically in women with current MDD. The association between depressive symptom severity and glutamine differed by CSA history. A significant interaction between baseline NAD /NADH ratio and clinical group predicted perceived stress at 6-month follow-up, adjusting for baseline perceived stress. Specifically, lower baseline NAD /NADH ratio predicted greater subsequent perceived stress in MDD without CSA, but not in the other clinical groups.

**Conclusions:** Our findings suggest that mPFC glutamatergic metabolites and redox balance show distinct patterns of association with CSA history, depression status, and subsequent perceived stress, highlighting neurochemical heterogeneity in MDD and molecular markers of risk.

## Introduction

Major depressive disorder (MDD) is among the most prevalent psychiatric disorders worldwide and a leading cause of disability (World Health Organization, 2017). Childhood maltreatment, including childhood sexual abuse (CSA), is a well-established risk factor for MDD (Li et al., 2016; Nelson et al., 2017). In women, CSA has been associated with more than a threefold increase in lifetime risk for MDD (Kendler et al., 2000), with elevated risk persisting into adulthood (Akinyemi et al., 2025).

Two interconnected neurobiological processes that may link early-life adversity to depression are glutamatergic signaling and cellular redox balance. Glutamatergic metabolism involves the glutamate– glutamine cycle, in which astrocytic glutamine synthetase converts glutamate to glutamine (J. V. Andersen et al., 2021). The medial prefrontal cortex (mPFC), a key region involved in stress regulation (Ulrich-Lai & Herman, 2009), has been implicated in glutamatergic alterations associated with depression. For example, in mice, disruption of astrocytic glutamine synthetase in the mPFC reduced glutamate and glutamine and induced depression-like behavior (Kang et al., 2026). Moreover, although acute stress rapidly increases extracellular glutamate release and excitatory transmission in rodents, particularly within the medial prefrontal cortex (mPFC; Moghaddam, 1993; Kaul et al., 2022), chronic or repeated stress has been associated with reduced prefrontal glutamatergic transmission, impaired astrocytic glutamate clearance and metabolism, and attenuated glutamate responses to subsequent stressors (e.g. Kaul et al., 2022; Treadway et al., 2025). Consistent with these observations, preclinical studies using proton magnetic resonance spectroscopy (MRS) and ¹H-[¹³C]-MRS have demonstrated that chronic stress induces depression-like behaviors accompanied by reduced prefrontal glutamate and glutamine concentrations, impaired neuronal and astroglial metabolism, and decreased glutamate– glutamine cycling (e.g. Veeraiah et al., 2014; Mishra et al., 2020). Recent mechanistic studies have further demonstrated that disruption of astrocytic glutamine synthetase activity or components of the astrocyte–neuron glutamate–glutamine cycle can induce depression-like behaviors, supporting a potentially causal contribution of impaired glutamatergic homeostasis (Kang et al., 2025, 2026). In humans, meta-analyses of MRS studies have identified reduced glutamate-plus-glutamine (Glx) concentrations in prefrontal and anterior cingulate regions in MDD, although changes in individual metabolites, including glutamine, vary across studies and brain regions (Tao et al., 2025; White et al., 2026). Convergent postmortem evidence implicates astrocyte pathology in MDD, including reduced astrocytic cell density or marker expression in some prefrontal regions and abnormalities in molecular systems responsible for glutamate uptake and metabolism (Miguel-Hidalgo et al., 2000; Liu et al., 2022; Lu et al., 2025; Nasu & Terunuma, 2026).

Cellular redox balance is closely linked to glutamatergic metabolism and can be assessed through the ratio of oxidized to reduced nicotinamide adenine dinucleotide (NAD /NADH), an indicator of cellular metabolic redox state (Xiao et al., 2018). These processes are interconnected through several metabolic pathways, including astrocytic glutamate uptake, glutamate–glutamine cycling, and mitochondrial energy metabolism (J. V. Andersen, 2025; McKenna et al., 2016). For example, astrocytic glutamate uptake has been shown to decrease the NAD /NADH ratio (Köhler et al., 2018). Moreover, chronic stress has been associated with alterations in mPFC NAD /NADH and depressive-like behavior in mice (Dong et al., 2025), and psychological stress more broadly alters mitochondrial function in animal models (Picard & McEwen, 2018). Thus, redox state dysregulation (specifically, a reduced NAD /NADH) might characterize MDD, particularly in the context of stress-related context, which likely affect anti-oxidant defenses, energy homeostasis as well as downstream effects on neuronal function and plasticity (Steullet et al., 2010). Together, these findings suggest that glutamatergic metabolism and cellular redox balance may contribute to stress-related neurobiological processes within the mPFC.

Indeed, both MDD and childhood maltreatment have been associated with initial evidence of alterations in glutamatergic metabolism and processes related to cellular redox balance. Specifically, MDD has characterized by increased peripheral markers of oxidative damage, including lipid peroxidation, protein oxidation, and oxidative DNA damage, alongside altered antioxidant defenses (Ait Tayeb et al., 2023). Similarly, childhood maltreatment has been linked to oxidative nucleoside damage (Eriksen et al., 2022), mitochondrial alterations (Zitkovsky et al., 2021), and reduced hippocampal glutamatergic metabolites in the context of mood disorders (Poletti et al., 2016). Moreover, adverse childhood experiences have been associated with lower glutamatergic metabolite levels in the mPFC of healthy adults (Duncan et al., 2015), suggesting that neurochemical alterations in this stress-regulatory region may extend beyond clinical populations. However, little is known about how CSA history and current versus remitted depression relate to mPFC glutamatergic metabolism and redox balance, particularly when these neurochemical measures are examined together.

Beyond clinical status, the severity and multiplicity of maltreatment exposure may contribute to neurochemical differences. Specifically, childhood maltreatment can influence brain development, with neurobiological alterations varying by exposure timing and type (Gee & Casey, 2015; Teicher et al., 2016). However, the associations of childhood maltreatment severity and multiplicity with mPFC glutamatergic metabolism and redox balance, and how these associations may vary with depressive symptom severity, remain understudied. Furthermore, although MRS-derived metabolites have been investigated as predictors of depression recurrence (Henigsberg et al., 2021), their potential to predict subsequent perceived stress remains largely unexplored.

To address these gaps, the present study investigated, we believe for the first time, mPFC glutamatergic metabolites and redox balance in women with varying histories of MDD and CSA. Specifically, we examined (1) group differences in glutamatergic metabolites and the NAD /NADH ratio across women with current MDD with and without CSA, remitted MDD with CSA, and healthy controls, hypothesizing that MDD and CSA, and possibly their co-occurrence would be associated with reduced glutamatergic metabolites and NAD /NADH ratio; (2) associations of these neurochemical measures with childhood maltreatment severity and multiplicity and depressive symptom severity, including potential differences across clinical groups; and (3) whether baseline neurochemical measures predicted perceived stress at 6-month follow-up.

## Methods and Materials

### Participants and study design

Women were recruited from the greater Boston area, primarily through social media advertisements and local college job boards, as part of a study investigating the functional and molecular effects of CSA occurring between ages 11–18 on depression (NCT04713722). We specifically selected the range of 11-18 for CSA occurrence, in light of findings that (1) PFC volume was reduced with ELS occurring at ages 14-16 years (S. L. Andersen et al., 2008) and similar findings that later (8-17 years), but not early (1 month - 7 years) childhood ELS was associated with reduced PFC volume (Baker et al., 2013); and (2) late adolescent (ages 12-17) exposure to abuse significantly increased MDD risk (Thornberry et al., 2001). Participants completed an initial clinical assessment, followed by an MRS session and a 6-month follow-up assessment. At the initial clinical visit, psychiatric diagnoses were established using the Structured Clinical Interview for DSM-5 (SCID-5-RV; First et al., 2015), and childhood maltreatment was assessed using the Maltreatment and Abuse Chronology of Exposure (MACE; Teicher & Parigger, 2015), with CSA histories confirmed through clinical interviews. Depressive symptom severity was measured using the Beck Depression Inventory-II (BDI-II; Beck et al., 1996). Perceived stress was assessed using the Perceived Stress Scale (PSS-14; Cohen et al., 1983) at the initial clinical visit and at 6-month follow-up.

The final sample for the current analyses comprised 71 women who underwent MRS (Table 1): healthy controls (HC; *n* = 27), current MDD without CSA (MDD/no CSA; *n* = 14), current MDD with CSA (MDD/CSA; *n* = 14), and remitted MDD with CSA (rMDD/CSA; *n* = 16). Three MDD/CSA participants were excluded from MACE analyses because of unreliable self-reported maltreatment histories, and two rMDD/CSA participants lacked glutamatergic metabolite data due to ¹H-MRS acquisition failure. Detailed exclusion criteria and variable-specific exclusions are provided in the *Supplement*. The study was approved by the Mass General Brigham and McLean Hospital Institutional Review Board (Protocol No. 2020P001470), and participants provided informed consent.

### MRS acquisition and quantification

Proton (¹H) and phosphorus (³¹P) MRS data were acquired from an overlapping region in the mPFC during a single session on a 4T MRI scanner (Varian/Agilent Whole Body DirectDrive™ scanner; Agilent Technologies, Palo Alto, CA) with a specially designed and dual-tuned half-helmet head coil. This RF probe consisted of a butterfly shaped ^1^H coil and a single-loop ^31^P coil (7-cm diameters). The detailed protocol of data acquisition and processing has been described in our previous publications (Cuklanz et al., 2026; Du et al., 2014; Kim et al., 2017). Briefly, the ^1^H MRS voxel was 25 × 20 × 25 mm, while the larger region around 60 × 60 × 40 mm covered by ^31^P RF coil was used to compensate for the intrinsically lower sensitivity of ^31^P MRS and the extremely low concentrations of NAD+ and NADH in the brain (both typically below 1mM), thereby improving signal-to noise ratio and enabling their reliable detection and qualification. Before spectroscopy data collection, localized shimming was performed using vendor-provided routines (3D-shimming) plus manual refinement if needed, achieving unsuppressed water linewidths < 12 and 25 Hz for ^1^H and ^31^P MRS, respectively.

Each scan (individual FID) was digitized, and frequency and phase drift as well as eddy current effects resulting from hardware variations or subject movement were corrected before summing all FIDs. Glutamate (Glu) and glutamine (Gln) were quantified using the SPECIAL sequence (repetition time/echo time = 3000/10 ms; 64 averages), with Glx calculated as their sum (Glu+Gln). Water suppression was implemented using VAPOR (Tkáč et al., 1999). Eight unsuppressed water signals were acquired for eddy-current correction and water-referenced quantification.

The NAD /NADH redox ratio, an indicator of cellular redox balance, was quantified using ³¹P-MRS with proton decoupling by a decoupling pulse train with WALTZ-16 (Cuklanz et al., 2026; Lu et al., 2016). A 300-µs hard pulse was used to excite the phosphate spins, and its flip angle (nominal 90°) was adjusted to achieve optimal nuclear magnetic resonance signal. The spectral width was 5000 Hz, repetition time was 3 s, and 640 averages were acquired. Total experimental time was approximately 75 minutes.

MRS scans were scheduled during the early follicular phase (days 1–7 of the menstrual cycle) whenever possible; however, 16 of the 64 participants with a recorded cycle day (25%) were scanned outside this window due to scheduling constraints.

All MRS data processing was blinded to diagnoses. Post-processing of FIDs includes zero filling, windowing functions, and Fourier transformations. MRS data processing was conducted by in-house software and LCModel (Provencher, 2001) with simulated basis. For ^1^H MRS, partial volume correction (Chen et al., 2019) was performed with gray matter, white matter, and CSF fractions obtained from T1- weighted images using SPM12 (Ashburner & Friston, 2005) and published T1/T2 values at 4T (Deelchand et al., 2012; Tkáč et al., 2009). For ^31^P MRS, initial 3 points of FIDs were truncated to remove the broad component. Nine metabolites (adenosine triphosphate, phosphocreatine, NAD , NADH, glycerophosphocholine, glycerophosphoethanolamine, inorganic phosphate, phosphocholine, and phosphoethanolamine) were included for basis simulation. Details of data processing – including the excellent test-retest reliability of the NAD+, NADH, and the ratio (Kim et al., 2017) – have been described in our previous publications (Cuklanz et al., 2026; Du et al., 2014; Kim et al., 2017; Yuksel et al., 2021).

### Clinical measures

Childhood maltreatment was assessed using the MACE, yielding two composite scores: *maltreatment multiplicity*, defined as the number of maltreatment subtypes exceeding published clinical-significance thresholds (range 0–10), and *total maltreatment severity*, calculated as the sum of severity scores across subtypes (range 0–100). Both measures reflected maltreatment exposure across childhood. MACE was scored across ages 11–18 to reflect the study’s CSA inclusion criteria. Perceived stress was assessed using the PSS (14 items) at baseline and 6-month follow-up (range: 0–56), with higher scores indicating greater perceived stress during the past month. Depressive symptoms were assessed using the BDI-II (range: 0–63), with higher scores indicating greater depressive symptom severity. Highlighting excellent internal reliability, Cronbach’s α in the present sample was 0.95 for the PSS-14 at baseline, 0.94 at 6-month follow-up, and 0.83 for the BDI-II among women with current MDD (see Supplementary Table S1 for values by group).

## Statistical Analyses

### Group differences in neurochemical measures

One-way analyses of variance (ANOVAs) were conducted in *R* to examine differences across the four groups (HC, MDD/no CSA, MDD/CSA, rMDD/CSA) in glutamate, glutamine, Glx, and NAD /NADH. A Bonferroni correction was applied to interpret findings, using a p value of 0.0125 (=0.05/4 metabolites). Post hoc pairwise comparisons were performed with Tukey adjustment.

### Associations of neurochemical measures with childhood maltreatment and depressive symptoms

General linear models (GLMs) were used to examine associations between neurochemical measures and measures of childhood maltreatment and depressive symptom severity. Separate models were fitted for each neurochemical measure (glutamate, glutamine, Glx, NAD /NADH), with the neurochemical measure as the dependent variable and clinical group, the relevant clinical measure (MACE or BDI-II), and their interaction as predictors. Analyses of childhood maltreatment were restricted to women with CSA histories (MDD/CSA and rMDD/CSA), whereas analyses of depressive symptom severity were restricted to women with current MDD (MDD/no CSA and MDD/CSA). Group was effect-coded, and continuous measures were mean-centered. BDI-II scores were log-transformed using ln(x + 1) prior to analysis due to their skewed distribution. Main effects and interactions were evaluated using Type III ANOVA *F*-tests applied to the fitted models. Significant interactions were followed by simple slope analyses.

### Neurochemical measures predicting subsequent perceived stress

GLMs were used to examine whether baseline neurochemical measures predicted perceived stress at 6-month follow-up among women in the three clinical groups (MDD/no CSA, MDD/CSA, and rMDD/CSA). Separate models were fitted for each neurochemical measure (glutamate, glutamine, Glx, and NAD /NADH ratio), with 6-month perceived stress as the dependent variable and clinical group, the neurochemical measure, and group-by-neurochemical measure interaction as predictors, adjusting for baseline perceived stress. Group was effect-coded, and continuous predictors were mean-centered. Main effects and interactions were evaluated using Type III ANOVA *F*-tests applied to the fitted models. Significant interactions were followed by simple slope analyses.

## Results

### Sample characteristics

The demographic and clinical characteristics are summarized in Table 1. No between-group differences in demographic variables were observed. The MDD groups did not differ in depression measures, and the CSA groups did not differ in maltreatment measures.

### Group differences in neurochemical measures

Representative ^1^H and ^31^P spectra are shown in Figure 1. Significant group differences emerged for glutamate (*F*(3, 64) = 5.25, *p* = .003; Figure 2A) and Glx (*F*(3, 65) = 5.62, *p* = .002; Figure 2B), with both findings surviving Bonferroni correction. Tukey-adjusted post hoc comparisons revealed lower glutamate and Glx levels in both the MDD/CSA and rMDD/CSA groups compared with HC. No significant group differences emerged for glutamine or the NAD /NADH ratio.

**Figure 1.**
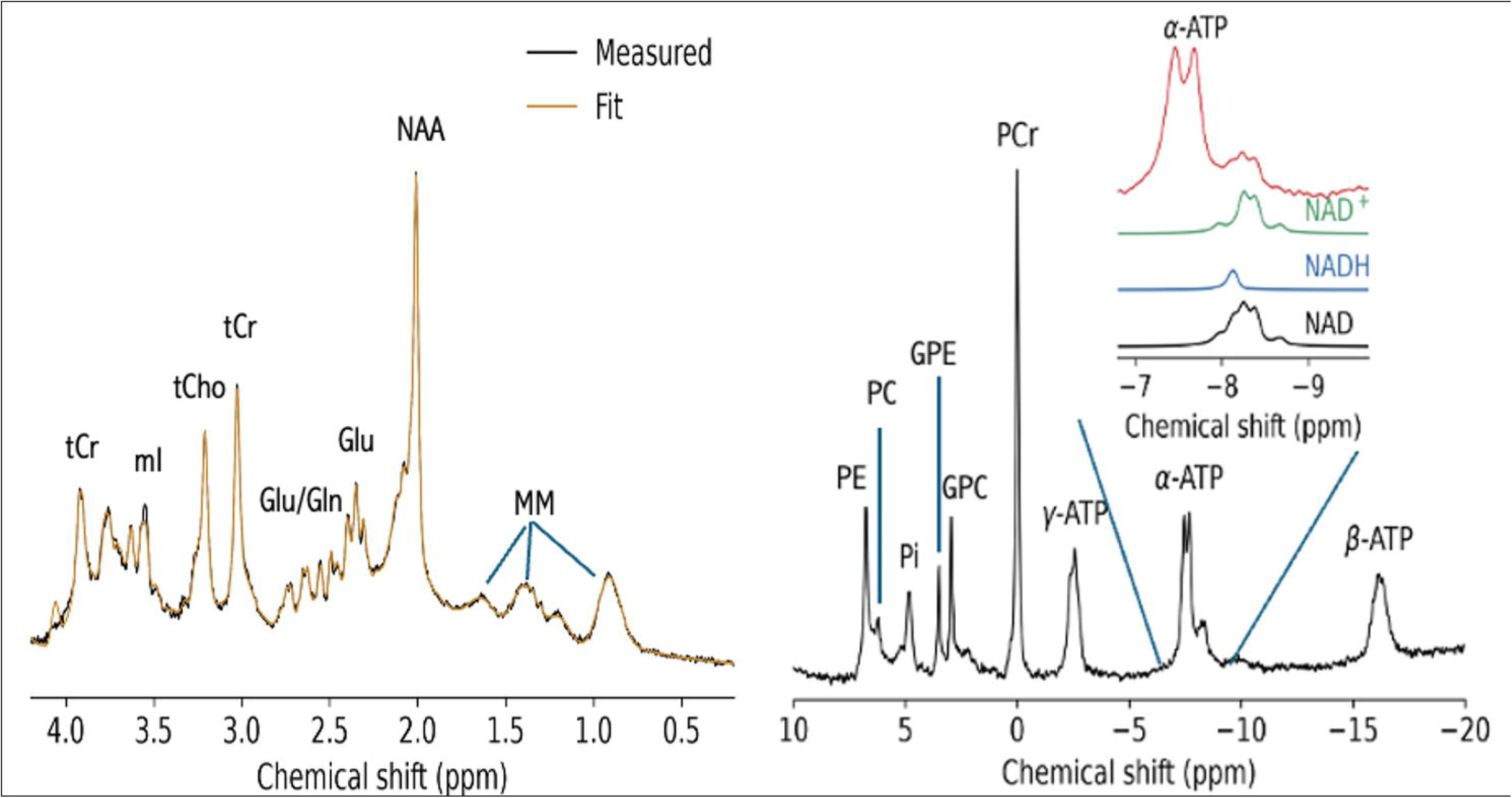
Representative in vivo ¹H and ³¹P MR spectra acquired from the medial prefrontal cortex on a 4 Tesla scanner. The ¹H spectrum shows the measured signal (black) and model fit (yellow). The ³¹P spectrum inset shows the NAD region, with the fitted NAD (green), NADH (blue), and total NAD (black) components. ATP, adenosine triphosphate; Gln, glutamine; Glu, glutamate; GPC, glycerophosphocholine; GPE, glycerophosphoethanolamine; mI, myo-inositol; MM, macromolecules; NAA, N-acetylaspartate; NAD /NADH, oxidized/reduced nicotinamide adenine dinucleotide (NAD); PC, phosphocholine; PCr, phosphocreatine; PE, phosphoethanolamine; Pi, inorganic phosphate; tCho, total choline; tCr, total creatine.

**Figure 2.**
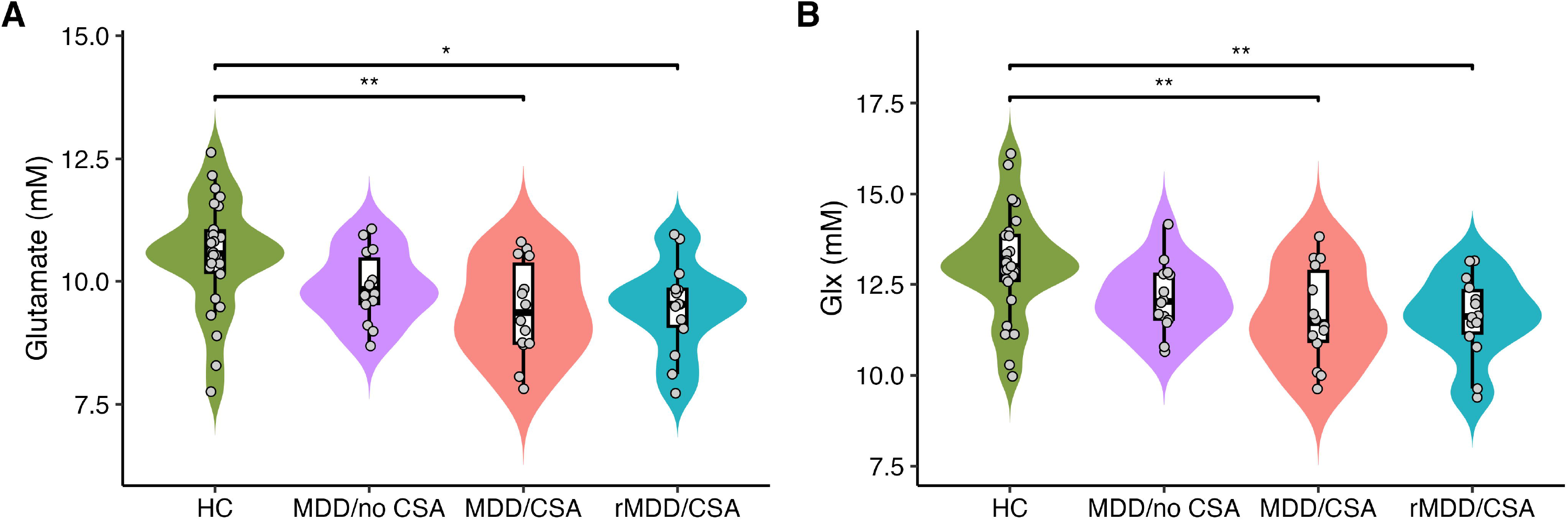
Group differences in mPFC glutamatergic metabolites. **(A)** Glutamate and **(B)** Glx (glutamate + glutamine) levels in the medial prefrontal cortex (mPFC) were significantly lower in women with MDD/CSA and rMDD/CSA than in HC. Violin plots show the distributions, boxes indicate medians and interquartile ranges, and points represent individual participants. Asterisks indicate significant Tukey-adjusted pairwise comparisons. CSA: childhood sexual abuse; HC: healthy controls; MDD: major depressive disorder; rMDD: remitted major depressive disorder.

### Associations of childhood maltreatment with glutamatergic metabolites

Among women with CSA histories (MDD/CSA + rMDD/CSA, *n* = 30), a significant interaction between maltreatment severity and depression status emerged for glutamine (*F*(1, 20) = 6.55, *p* = .019; Figure 3A). Simple slope analyses indicated a significant negative association between maltreatment severity and glutamine in MDD/CSA (*t*(20) = −2.29, *p* = .032), but not rMDD/CSA (*t*(20) = 1.14, *p* = .267). Significant interactions between maltreatment multiplicity and depression status were observed for glutamate (*F*(1, 21) = 4.66, *p* = .043; Figure 3B) and Glx (*F*(1, 21) = 4.58, *p* = .044; Figure 3C). Greater maltreatment multiplicity related to lower glutamate (*t*(21) = −2.65, *p* = .015) and Glx (*t*(21) = −2.56, *p* = .018) in MDD/CSA, but not in rMDD/CSA (glutamate: *t*(21) = −0.42, *p* = .682; Glx: *t*(21) = −0.29, *p* = .778).

**Figure 3.**
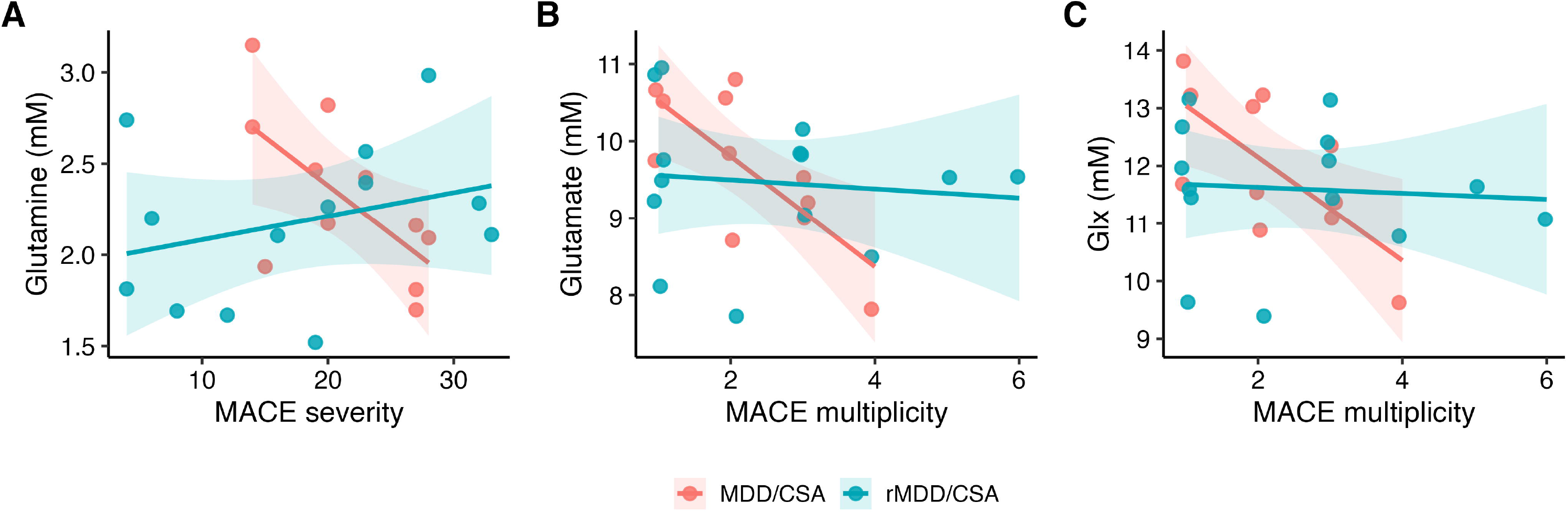
Associations between mPFC glutamatergic metabolites and childhood maltreatment differ by depression status. Among women with CSA histories, depression status significantly moderated the associations between **(A)** maltreatment severity and medial prefrontal cortex (mPFC) glutamine, **(B)** maltreatment multiplicity and mPFC glutamate, and **(C)** maltreatment multiplicity and mPFC Glx (glutamate + glutamine). Greater maltreatment severity and multiplicity were associated with lower levels of the respective glutamatergic metabolites in women with current MDD, whereas these associations were not significant in women with remitted MDD. Shaded regions represent 95% confidence intervals around the fitted regression lines. CSA: childhood sexual abuse; MACE: Maltreatment and Abuse Chronology of Exposure; MDD: major depressive disorder; rMDD: remitted major depressive disorder.

### Associations of depressive symptom severity with glutamatergic metabolites

Among women with current MDD (MDD/no CSA + MDD/CSA, *n* = 28), a significant interaction between depressive symptom severity and CSA history was observed for glutamine (*F*(1, 24) = 4.28, *p* =.049; Figure 4). Simple slope analyses revealed a trend-level negative association between depressive symptom severity and glutamine in MDD/no CSA (*t*(24) = −1.94, *p* = .064), but not MDD/CSA (*t*(24) = 0.87, *p* = .395).

**Figure 4.**
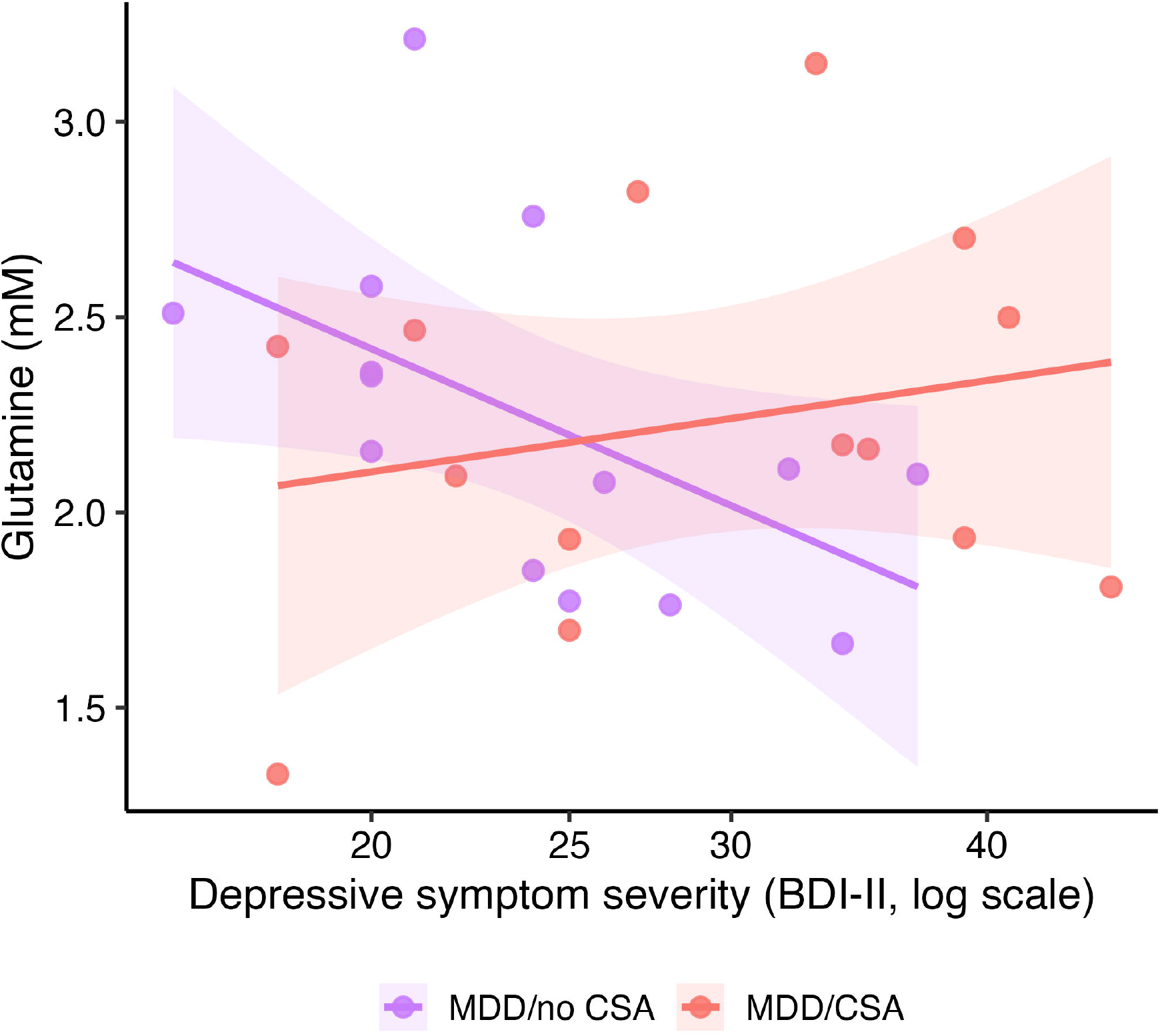
Associations between mPFC glutamine and depression severity differ by CSA history. Among women with current MDD, the association between depressive symptom severity (BDI-II scores) and medial prefrontal cortex (mPFC) glutamine differed significantly by CSA history. A trend-level negative association was observed in women without CSA, whereas no significant association was observed in women with CSA. BDI-II scores were log-transformed using ln(x + 1) and are displayed on the transformed scale. Shaded regions represent 95% confidence intervals around the fitted regression lines. BDI-II, Beck Depression Inventory-II; CSA, childhood sexual abuse; MDD, major depressive disorder.

### Redox balance predicting subsequent perceived stress

Among women in the three clinical groups (MDD/no CSA + MDD/CSA + rMDD/CSA: *n* = 44), a significant interaction between baseline NAD /NADH ratio and clinical group was observed in predicting perceived stress at 6-month follow-up, adjusting for baseline perceived stress (*F*(2, 27) = 8.50, *p* = .001; Figure 5). Simple slope analyses revealed that a lower baseline NAD /NADH ratio predicted greater perceived stress at 6-month follow-up in MDD/no CSA (*t*(27) = −4.29, *p* < .001), but not in MDD/CSA (*t*(27) = −0.39, *p* = .698) or rMDD/CSA (*t*(27) = 1.31, *p* = .201).

**Figure 5.**
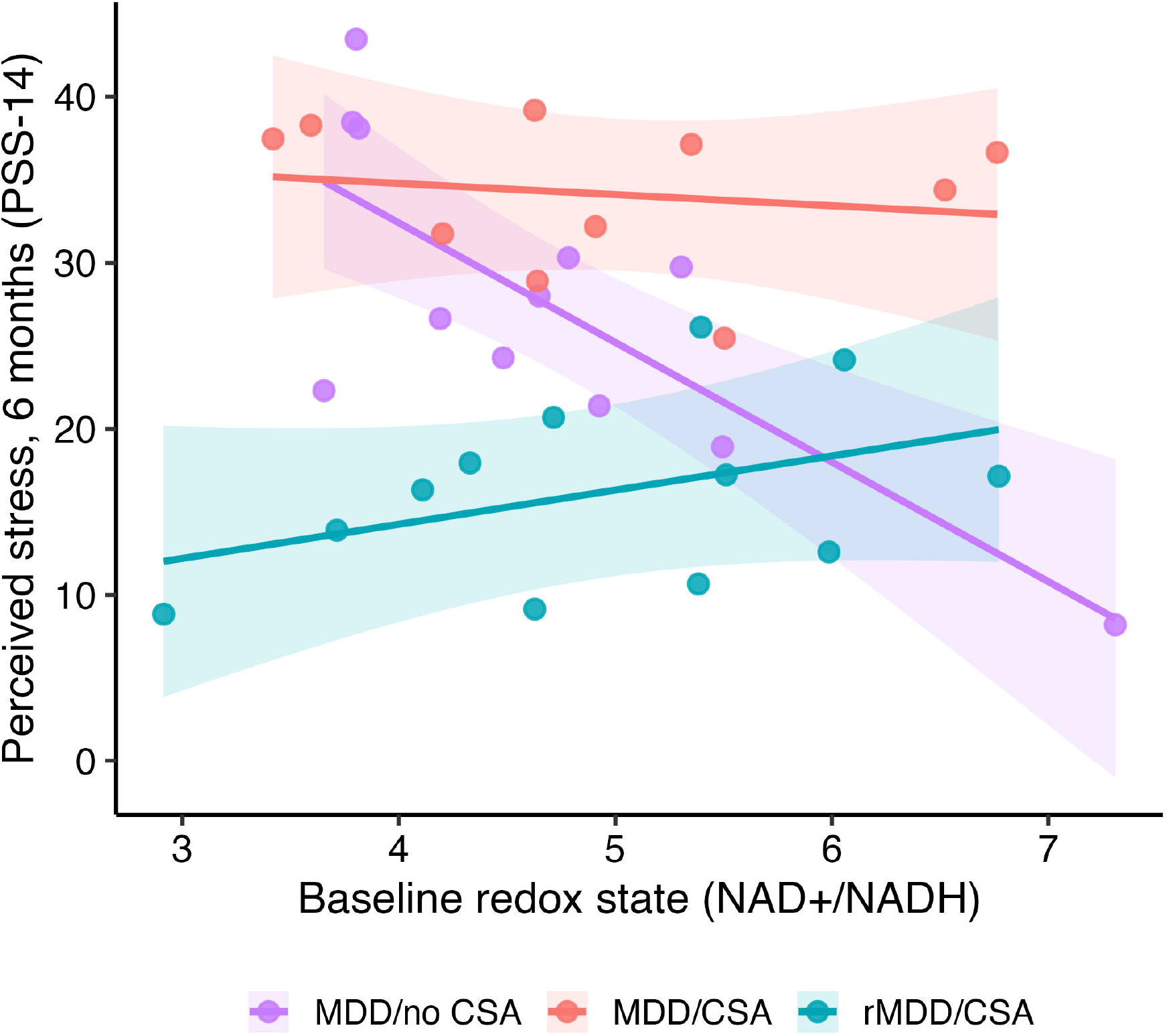
Associations between baseline mPFC NAD /NADH ratio and subsequent perceived stress differ by clinical group. Among women with current or remitted MDD, the association between baseline mPFC NAD /NADH ratio and perceived stress at 6-month follow-up differed significantly across clinical groups, adjusting for baseline perceived stress. A lower baseline NAD /NADH ratio predicted greater subsequent perceived stress in women with current MDD without CSA, whereas no significant associations were observed in either CSA-exposed group. Points represent 6-month PSS-14 scores adjusted for baseline PSS-14, and lines represent model-predicted values at the mean baseline PSS-14 score. Shaded regions indicate 95% confidence intervals around the fitted regression lines. CSA, childhood sexual abuse; MDD, major depressive disorder; NAD /NADH, ratio of oxidized to reduced nicotinamide adenine dinucleotide; PSS-14, 14-item Perceived Stress Scale; rMDD, remitted major depressive disorder.

## Discussion

The overarching goal of the current study was to test, we believe for the first time, mPFC glutamatergic metabolites and redox balance in relation to childhood maltreatment, depression, and perceived stress among unmedicated women differing in CSA history and depression status. Three main findings emerged. First, glutamate and Glx levels were lower in women with lifetime MDD and CSA histories compared with HC, whereas women with MDD without CSA did not differ significantly from HC. Second, associations between glutamatergic metabolites and clinical characteristics varied according to CSA history and depression status. Specifically, greater maltreatment severity and multiplicity were associated with lower glutamatergic metabolite levels among women with current MDD and CSA histories, but not among those with remitted MDD. Additionally, among women with current MDD, the association between depressive symptom severity and glutamine differed by CSA history. Third, a lower baseline NAD /NADH ratio predicted greater perceived stress 6 months later among women with current MDD without CSA, but not among women with CSA histories. Together, these findings highlight the importance of considering both childhood maltreatment and depression status when examining mPFC neurochemistry and its relationship to clinical symptoms and subsequent stress vulnerability.

The lower mPFC glutamate and Glx levels observed in both CSA-exposed groups, including women with remitted MDD, are consistent with previous studies linking childhood adversity to lower mPFC glutamate in healthy adults (Duncan et al., 2015) and severe trauma during adolescence to lower ventromedial prefrontal glutamate (Ousdal et al., 2019). Although a meta-analysis identified reduced medial frontal Glx in MDD, substantial heterogeneity across studies was evident (Moriguchi et al., 2019). Studies of remitted depression have likewise reported differing glutamatergic findings, including lower glutamate in the ventromedial (Portella et al., 2011) and dorsolateral (Ritter et al., 2022) prefrontal cortex, and normal glutamate in the anterior cingulate cortex (Taylor et al., 2009). Together, these findings raise the possibility that maltreatment history contributes to heterogeneity in glutamatergic metabolite levels in MDD, alongside differences in brain regions, measurement methods, and clinical characteristics. However, the absence of a CSA-exposed group without lifetime MDD precludes attributing the observed group differences specifically to maltreatment.

Beyond these group-level differences, associations with maltreatment severity and depressive symptoms provided further insight into the clinical correlates of glutamatergic metabolism. Among CSA- exposed women, greater maltreatment severity and multiplicity were associated with lower glutamatergic metabolite levels during current, but not remitted, MDD. Among women with current MDD, the association between depressive symptom severity and glutamine differed by CSA history, with a trend-level negative association in those without CSA. Prior MRS studies of childhood adversity in depressed samples have yielded mixed findings: adversity has been associated with lower hippocampal Glx/creatine ratio in patients with mood disorders (Poletti et al., 2016), but with higher occipital glutamine and glutamate–glutamine cycling in adults with MDD (Averill et al., 2020) and higher anterior cingulate Glx in depressed adolescents (Sonmez et al., 2021). None of these studies distinguished current from remitted depression. Because maltreatment-related associations in the present sample differed between current and remitted MDD, depression status may help explain some of this heterogeneity. These findings are consistent with the proposed distinction between maltreatment-exposed and non-exposed forms of depression, which may differ in underlying neurobiology (Teicher & Samson, 2013). Notably, although both CSA-exposed groups were characterized by lower glutamate and Glx levels than HC, associations with maltreatment severity and multiplicity emerged only in current MDD, suggesting that depression status may influence how childhood adversity relates to glutamatergic metabolism.

The prospective analysis revealed that the association between baseline mPFC NAD /NADH ratio and perceived stress 6 months later differed across clinical groups. Specifically, a lower baseline NAD /NADH ratio predicted greater subsequent perceived stress among women with current MDD without CSA, after adjusting for baseline perceived stress, whereas no significant associations were observed in either CSA-exposed group. Although group differences in NAD /NADH did not emerge, the direction of this association, in which a lower NAD /NADH ratio predicted greater subsequent perceived stress, aligns with previous ^31^P MRS studies reporting reduced NAD /NADH ratios in schizophrenia (Kim et al., 2017) and first-episode psychosis, including bipolar disorder with psychotic features (Chouinard et al., 2017). However, little research has examined whether redox balance predicts subsequent stress-related outcomes in the context of depression. Previous longitudinal MRS research has linked neurochemical measures to depression recurrence, with Henigsberg et al. (2021) reporting that higher amygdala-region choline-containing metabolite levels during recovery were associated with a lower risk of subsequent depressive episodes, leaving the prospective clinical relevance of redox balance in depression largely unexplored. Our findings extend this work by identifying a prospective association between mPFC redox balance and perceived stress that depends on CSA history and depression status. To our knowledge, ours is among the first studies to measure mPFC redox balance alongside glutamatergic metabolites in the same voxel in women with MDD and childhood maltreatment histories. Given the role of NAD /NADH in energy metabolism, mitochondrial function, inflammation and oxidative stress (Xiao et al., 2018), this approach provides a unique window into cellular processes that have been difficult to study in the living human brain. Although these findings suggest that baseline NAD /NADH ratio may provide information beyond baseline stress levels, its incremental predictive value beyond other clinical characteristics and its generalizability require further investigation.

Several limitations warrant consideration. First, although the multi-modal assessment, unmedicated status, and deep phenotyping of the parent study is a strength, subgroup sizes were modest, particularly for interaction analyses, and, beyond the main analyses comparing groups in the four molecular markers, no correction for multiple testing was applied given the exploratory nature of the study. Second, menstrual cycle phase was not uniformly controlled; although scanning was targeted to the early follicular phase when feasible, variability in scan timing may have influenced neurochemical measurements. Third, the absence of a CSA-exposed group without lifetime MDD limits the ability to disentangle associations related to maltreatment history, depression, or their combination. Fourth, neurochemical measures were obtained at a single time point, limiting conclusions about temporal relationships between neurochemistry and clinical characteristics, while the prospective association with perceived stress requires independent validation. Fifth, neurochemical measures were quantified within a single mPFC voxel, limiting the ability to determine whether the findings vary across regions. Additionally, proton MRS cannot distinguish neuronal from astrocytic or intracellular from extracellular glutamate and glutamine. Thus, larger longitudinal studies are needed to replicate these findings and clarify the underlying mechanisms.

Finally, the separate quantification of Glu and Gln presents methodological challenges that warrant consideration when interpreting our glutamatergic findings. To separately quantify Glu and Gln, we implemented a SPECIAL sequence with adiabatic RF pulses and an ultrashort echo time (TE = 10 ms) at 4T. Compared with STEAM, SPECIAL provides greater signal intensity, improving signal-to-noise ratio while maintaining a short TE. In eight healthy volunteers, LCModel fitting yielded low Cramér–Rao lower bounds (2.6%, 8.7%, and 10.5% for Glu, Gln, and GSH, respectively) and low fitting correlations between Glu and Gln (0.05), Glu and GSH (−0.11), and Gln and GSH (−0.15), supporting the feasibility of separately quantifying these metabolites. Estimated metabolite concentrations were also consistent with prior literature. Nevertheless, Glu and Gln exhibit overlapping spectral resonances and are subject to J-coupling, introducing potential uncertainty in their individual estimates. We therefore also examined Glx (Glu + Gln) as a complementary measure that is less dependent on distinguishing the contributions of these overlapping metabolites.

## Conclusions

The present findings highlight the importance of childhood maltreatment history and depression status in understanding mPFC glutamatergic metabolites and redox balance in MDD. Rather than uniformly characterizing MDD, these neurochemical measures showed distinct patterns of association with maltreatment history, depressive symptoms, and subsequent perceived stress. Larger longitudinal studies are needed to determine whether integrating maltreatment history and neurochemical measures can improve our understanding of neurobiological heterogeneity and clinical trajectories in MDD.

## Data availability

De-identified, subject-level spectroscopy data are subject to institutional policies, IRB requirements, and study-specific data-use agreements. Data processing software and analysis code will be made available through an appropriate repository, to the extent permitted by applicable institutional policies, ethical approvals, and data use agreements.

## Acknowledgements

This study was supported by the National Institute of Mental Health (NIMH) R01 MH095809 awarded to DAP and FD.

## References

1. Ait Tayeb, A. E. K., Poinsignon, V., Chappell, K., Bouligand, J., Becquemont, L., & Verstuyft, C. (2023). Major depressive disorder and oxidative stress: A review of peripheral and genetic biomarkers according to clinical characteristics and disease stages. Antioxidants, 12(4), 942. 10.3390/antiox12040942

2. Akinyemi, O., Ogundare, T., Fasokun, M., Ogunyankin, F., Ugochukwu, N., Ajisafe, W., Ikugbayigbe, S., Eze, O., Hughes, K., & Michael, M. (2025). The long-term impact of childhood sexual assault on depression and self-reported mental and physical health. Frontiers in Psychiatry, 15, 1528914. 10.3389/fpsyt.2024.1528914

3. Andersen, J. V. (2025). The glutamate/GABA-glutamine cycle: Insights, updates, and advances. Journal of Neurochemistry, 169(3), e70029. 10.1111/jnc.70029

4. Andersen, J. V., Markussen, K. H., Jakobsen, E., Schousboe, A., Waagepetersen, H. S., Rosenberg, P. A., & Aldana, B. I. (2021). Glutamate metabolism and recycling at the excitatory synapse in health and neurodegeneration. Neuropharmacology, 196, 108719. 10.1016/j.neuropharm.2021.108719

5. Andersen, S. L., Tomada, A., Vincow, E. S., Valente, E., Polcari, A., & Teicher, M. H. (2008). Preliminary evidence for sensitive periods in the effect of childhood sexual abuse on regional brain development. Journal of Neuropsychiatry and Clinical Neurosciences, 20(3), 292–301.

6. Ashburner, J., & Friston, K. J. (2005). Unified segmentation. NeuroImage, 26(3), 839–851.

7. Averill, L. A., Abdallah, C. G., Fenton, L. R., Fasula, M. K., Jiang, L., Rothman, D. L., Mason, G. F., & Sanacora, G. (2020). Early life stress and glutamate neurotransmission in major depressive disorder. European Neuropsychopharmacology, 35, 71–80.

8. Baker, L. M., Williams, L. M., Korgaonkar, M. S., Cohen, R. A., Heaps, J. M., & Paul, R. H. (2013). Impact of early vs. late childhood early life stress on brain morphometrics. Brain Imaging and Behavior, 7(2), 196–203.

9. Beck, A. T., Steer, R. A., & Brown, G. K. (1996). Manual for the Beck Depression Inventory-II. Psychological Corporation.

10. Chen, X., Fan, X., Hu, Y., Zuo, C., Whitfield-Gabrieli, S., Holt, D., Gong, Q., Yang, Y., Pizzagalli, D. A., Du, F., & Öngür, D. (2019). Regional GABA concentrations modulate inter-network resting-state functional connectivity. Cerebral Cortex, 29(4), 1607–1618.

11. Chouinard, V.-A., Kim, S.-Y., Valeri, L., Yuksel, C., Ryan, K. P., Chouinard, G., Cohen, B. M., Du, F., & Öngür, D. (2017). Brain bioenergetics and redox state measured by ³¹P magnetic resonance spectroscopy in unaffected siblings of patients with psychotic disorders. Schizophrenia Research, 187, 11–16.

12. Cohen, S., Kamarck, T., & Mermelstein, R. (1983). A global measure of perceived stress. Journal of Health and Social Behavior, 24(4), 385–396.

13. Cooper, J. A., Nuutinen, M. R., Lawlor, V. M., DeVries, B. A. M., Barrick, E. M., Hossein, S., Cole, D. J., Leonard, C. V., Hahn, E. C., Teer, A. P., Shields, G. S., Slavich, G. M., Öngür, D., Jensen, J. E., Du, F., Pizzagalli, D. A., & Treadway, M. T. (2021). Reduced adaptation of glutamatergic stress response is associated with pessimistic expectations in depression. Nature Communications, 12, 3166. 10.1038/s41467-021-23284-9

14. Cuklanz, K. W., Stein, A., Chouinard, V.-A., Öngür, D., & Du, F. (2026). Redox therapy for neuropsychiatric disorders: Molecular mechanisms and biomarker development. Science Advances, 12(8), eaea9014. 10.1126/sciadv.aea9014

15. Deelchand, D. K., Henry, P.-G., Uğurbil, K., & Marjańska, M. (2012). Measurement of transverse relaxation times of J-coupled metabolites in the human visual cortex at 4 T. Magnetic Resonance in Medicine, 67(4), 891–897.

16. Dong, W., Liu, D., Fu, S., Zhang, J., Chen, X., & Huang, S. (2025). Mitochondrial CISD1 modulates microglial metabolic reprogramming to drive stress susceptibility in mice. *Advanced Science*, e08957. 10.1002/advs.202508957

17. Du, F., Cooper, A. J., Thida, T., Sehovic, S., Lukas, S. E., Cohen, B. M., Zhang, X., & Öngür, D. (2014). In vivo evidence for cerebral bioenergetic abnormalities in schizophrenia measured using ³¹P magnetization transfer spectroscopy. JAMA Psychiatry, 71(1), 19–27.

18. Duncan, N. W., Hayes, D. J., Wiebking, C., Tiret, B., Pietruska, K., Chen, D. Q., Rainville, P., Marjańska, M., Ayad, O., Doyon, J., Hodaie, M., & Northoff, G. (2015). Negative childhood experiences alter a prefrontal-insular-motor cortical network in healthy adults: A preliminary multimodal rsfMRI- fMRI-MRS-dMRI study. Human Brain Mapping, 36(11), 4622–4637.

19. Eriksen, J. K. D., Coello, K., Stanislaus, S., Kjærstad, H. L., Sletved, K. S. O., McIntyre, R. S., Faurholt-Jepsen, M., Miskowiak, K. K., Poulsen, H. E., Kessing, L. V., & Vinberg, M. (2022). Associations between childhood maltreatment and oxidative nucleoside damage in affective disorders. European Psychiatry, 65(1), e46. 10.1192/j.eurpsy.2022.2300

20. First, M. B., Williams, J. B. W., Karg, R. S., & Spitzer, R. L. (2015). Structured clinical interview for DSM-5—Research version (SCID-5 for DSM-5, research version; SCID-5-RV). American Psychiatric Association.

21. Gee, D. G., & Casey, B. J. (2015). The impact of developmental timing for stress and recovery. Neurobiology of Stress, 1, 184–194.

22. Henigsberg, N., Savić, A., Radoš, M., Radoš, M., Šarac, H., Šečić, A., Bajs Janović, M., Foro, T., Ozretić, D., Erdeljić Turk, V., Hrabač, P., & Kalember, P. (2021). Choline elevation in amygdala region at recovery indicates longer survival without depressive episode: A magnetic resonance spectroscopy study. Psychopharmacology, 238(5), 1303–1314.

23. Kang JS, Baek JH, Park H, Rehman NU, Chung HJ, Lee DK, et al. Disrupted astrocyte-neuron glutamine-glutamate cycling in the medial prefrontal cortex contributes to depression-like behaviors. Int J Biol Sci. 2026;22(2):553–565.

24. Kang JS, Kim H, Baek JH, Song M, Park H, Jeong W, et al. Activation of glutamine synthetase (GS) as a new strategy for the treatment of major depressive disorder and other GS-related diseases. Acta Pharmacol Sin. 2025;46(4):880–891.

25. Kaul D, Schwab SG, Mechawar N, Ooi L, Matosin N. Alterations in astrocytic regulation of excitation and inhibition by stress exposure and in severe psychopathology. J Neurosci. 2022;42(36):6823–6834.

26. Kendler, K. S., Bulik, C. M., Silberg, J., Hettema, J. M., Myers, J., & Prescott, C. A. (2000). Childhood sexual abuse and adult psychiatric and substance use disorders in women: An epidemiological and cotwin control analysis. Archives of General Psychiatry, 57(10), 953–959.

27. Kim, S.-Y., Cohen, B. M., Chen, X., Lukas, S. E., Shinn, A. K., Yuksel, A. C., Li, T., Du, F., & Öngür, D. (2017). Redox dysregulation in schizophrenia revealed by in vivo NAD /NADH measurement. Schizophrenia Bulletin, 43(1), 197–204.

28. Köhler, S., Winkler, U., Sicker, M., & Hirrlinger, J. (2018). NBCe1 mediates the regulation of the NADH/NAD redox state in cortical astrocytes by neuronal signals. Glia, 66(10), 2233–2245.

29. Li, M., D’Arcy, C., & Meng, X. (2016). Maltreatment in childhood substantially increases the risk of adult depression and anxiety in prospective cohort studies: Systematic review, meta-analysis, and proportional attributable fractions. Psychological Medicine, 46(4), 717–730.

30. Liu S-H, Du Y, Chen L, Cheng Y. Glial cell abnormalities in major psychiatric diseases: A systematic review of postmortem brain studies. Mol Neurobiol. 2022;59(3):1665–1692.

31. Lu C-L, Ren J, Cao X. An astroglial basis of major depressive disorder: Molecular, cellular, and circuit features. Biol Psychiatry. 2025;97(3):217–226.

32. Lu, M., Zhu, X.-H., & Chen, W. (2016). In vivo ³¹P MRS assessment of intracellular NAD metabolites and NAD /NADH redox state in human brain at 4 Tesla. NMR in Biomedicine, 29(7), 1010– 1017.

33. McKenna, M. C., Stridh, M. H., McNair, L. F., Sonnewald, U., Waagepetersen, H. S., & Schousboe, A. (2016). Glutamate oxidation in astrocytes: Roles of glutamate dehydrogenase and aminotransferases. Journal of Neuroscience Research, 94(12), 1561–1571.

34. Miguel-Hidalgo, J. J., Baucom, C., Dilley, G., Overholser, J. C., Meltzer, H. Y., Stockmeier, C. A., & Rajkowska, G. (2000). Glial fibrillary acidic protein immunoreactivity in the prefrontal cortex distinguishes younger from older adults in major depressive disorder. Biological Psychiatry, 48(8), 861–873.

35. Mishra P, Adusumilli M, Deolal P, Mason GF, Kumar A, Patel AB. Impaired neuronal and astroglial metabolic activity in chronic unpredictable mild stress model of depression: Reversal of behavioral and metabolic deficit with lanicemine. Neurochem Int. 2020:137:104750. doi: 10.1016/j.neuint.2020.104750.

36. Moghaddam, B. (1993). Stress preferentially increases extraneuronal levels of excitatory amino acids in the prefrontal cortex: Comparison to hippocampus and basal ganglia. Journal of Neurochemistry, 60(5), 1650–1657.

37. Moriguchi, S., Takamiya, A., Noda, Y., Horita, N., Wada, M., Tsugawa, S., Plitman, E., Sano, Y., Tarumi, R., ElSalhy, M., Katayama, N., Ogyu, K., Miyazaki, T., Kishimoto, T., Graff-Guerrero, A., Meyer, J. H., Blumberger, D. M., Daskalakis, Z. J., Mimura, M., & Nakajima, S. (2019). Glutamatergic neurometabolite levels in major depressive disorder: A systematic review and meta-analysis of proton magnetic resonance spectroscopy studies. Molecular Psychiatry, 24(7), 952–964.

38. Nasu Y, Terunuma M. From metabolism to mood regulation: astrocytes as a driver of depression. Front Cell Neurosci. 2026:20:1776649. doi: 10.3389/fncel.2026.1776649.

39. Nelson, J., Klumparendt, A., Doebler, P., & Ehring, T. (2017). Childhood maltreatment and characteristics of adult depression: Meta-analysis. British Journal of Psychiatry, 210(2), 96–104.

40. Ousdal, O. T., Milde, A. M., Craven, A. R., Ersland, L., Endestad, T., Melinder, A., Huys, Q. J., & Hugdahl, K. (2019). Prefrontal glutamate levels predict altered amygdala–prefrontal connectivity in traumatized youths. Psychological Medicine, 49(11), 1822–1830.

41. Picard, M., & McEwen, B. S. (2018). Psychological stress and mitochondria: A systematic review. Psychosomatic Medicine, 80(2), 141–153.

42. Poletti, S., Locatelli, C., Falini, A., Colombo, C., & Benedetti, F. (2016). Adverse childhood experiences associate to reduced glutamate levels in the hippocampus of patients affected by mood disorders. Progress in Neuro-Psychopharmacology & Biological Psychiatry, 71, 117–122.

43. Portella, M. J., de Diego-Adeliño, J., Gómez-Ansón, B., Morgan-Ferrando, R., Vives, Y., Puigdemont, D., Pérez-Egea, R., Ruscalleda, J., Álvarez, E., & Pérez, V. (2011). Ventromedial prefrontal spectroscopic abnormalities over the course of depression: A comparison among first episode, remitted recurrent and chronic patients. Journal of Psychiatric Research, 45(4), 427–434.

44. Provencher, S. W. (2001). Automatic quantitation of localized in vivo ¹H spectra with LCModel. NMR in Biomedicine, 14(4), 260–264.

45. Ritter, C., Buchmann, A., Müller, S. T., Volleberg, M., Haynes, M., Ghisleni, C., Noeske, R., Tuura, R., & Hasler, G. (2022). Evaluation of prefrontal γ-aminobutyric acid and glutamate levels in individuals with major depressive disorder using proton magnetic resonance spectroscopy. JAMA Psychiatry, 79(12), 1209–1216.

46. Shirayama, Y., Takahashi, M., Osone, F., Hara, A., & Okubo, T. (2017). Myo-inositol, glutamate, and glutamine in the prefrontal cortex, hippocampus, and amygdala in major depression. Biological Psychiatry: Cognitive Neuroscience and Neuroimaging, 2(2), 196–204.

47. Sonmez, A. I., Lewis, C. P., Port, J. D., Athreya, A. P., Choi, D.-S., Zaccariello, M. J., Shekunov, J., Blacker, C. J., & Croarkin, P. E. (2021). A pilot spectroscopy study of adversity in adolescents. Biomarkers in Neuropsychiatry, 5, 100043. 10.1016/j.bionps.2021.100043

48. Steullet, P., Cabungcal, J.-H., Kulak, A., Kraftsik, R., Chen, Y., Dalton, T. P., Cuenod, M., & Do, K. Q. (2010). Redox dysregulation affects the ventral but not dorsal hippocampus: Impairment of parvalbumin neurons, gamma oscillations, and related behaviors. Journal of Neuroscience, 30(7), 2547–2558.

49. Tao S, Deng R, Wei M, Huang Y, Sun H, Yang S, Li S, Xiao C, Li M. A meta-analysis of magnetic resonance spectroscopy studies on glutamatergic neurometabolite levels in major depressive disorder. Depress Anxiety. 2025:2025:5180077. doi: 10.1155/da/5180077.

50. Taylor, M. J., Selvaraj, S., Norbury, R., Jezzard, P., & Cowen, P. J. (2009). Normal glutamate but elevated myo-inositol in anterior cingulate cortex in recovered depressed patients. Journal of Affective Disorders, 119(1–3), 186–189.

51. Teicher, M. H., & Parigger, A. (2015). The ’Maltreatment and Abuse Chronology of Exposure’ (MACE) scale for the retrospective assessment of abuse and neglect during development. PLoS ONE, 10(2), e0117423. 10.1371/journal.pone.0117423

52. Teicher, M. H., & Samson, J. A. (2013). Childhood maltreatment and psychopathology: A case for ecophenotypic variants as clinically and neurobiologically distinct subtypes. American Journal of Psychiatry, 170(10), 1114–1133.

53. Teicher, M. H., Samson, J. A., Anderson, C. M., & Ohashi, K. (2016). The effects of childhood maltreatment on brain structure, function and connectivity. Nature Reviews Neuroscience, 17(10), 652–666.

54. Thornberry, T. P., Ireland, T. O., & Smith, C. A. (2001). The importance of timing: The varying impact of childhood and adolescent maltreatment on multiple problem outcomes. Development and Psychopathology, 13(4), 957–979.

55. Tkáč, I., Öz, G., Adriany, G., Uğurbil, K., & Gruetter, R. (2009). In vivo ¹H NMR spectroscopy of the human brain at high magnetic fields: Metabolite quantification at 4T vs. 7T. Magnetic Resonance in Medicine, 62(4), 868–879.

56. Tkáč, I., Starčuk, Z., Choi, I.-Y., & Gruetter, R. (1999). In vivo ¹H NMR spectroscopy of rat brain at 1 ms echo time. Magnetic Resonance in Medicine, 41(4), 649–656.

57. Ulrich-Lai, Y. M., & Herman, J. P. (2009). Neural regulation of endocrine and autonomic stress responses. Nature Reviews Neuroscience, 10(6), 397–409.

58. Veeraiah, P., Noronha, J. M., Maitra, S., Bagga, P., Khandelwal, N., Chakravarty, S., Kumar, A., & Patel, A. B. (2014). Dysfunctional glutamatergic and γ-aminobutyric acidergic activities in prefrontal cortex of mice in social defeat model of depression. Biological Psychiatry, 76(3), 231–238.

59. White AJ, Robinson JL, Wilson AE, Reid MA. Neurometabolics of the anterior cingulate cortex (ACC) in major depressive disorder (MDD): A systematic review and meta-analysis. Neuroimage Clin. 2026:49:103959. doi: 10.1016/j.nicl.2026.103959.

60. World Health Organization. (2017). Depression and other common mental disorders: Global health estimates. World Health Organization.

61. Xiao, W., Wang, R.-S., Handy, D. E., & Loscalzo, J. (2018). NAD(H) and NADP(H) redox couples and cellular energy metabolism. Antioxidants & Redox Signaling, 28(3), 251–272.

62. Yuksel, C., Chen, X., Chouinard, V.-A., Nickerson, L. D., Gardner, M., Cohen, T., Öngür, D., & Du, F. (2021). Abnormal brain bioenergetics in first-episode psychosis. Schizophrenia Bulletin Open, 2(1), Article sgaa073. 10.1093/schizbullopen/sgaa073

63. Zitkovsky, E. K., Daniels, T. E., & Tyrka, A. R. (2021). Mitochondria and early-life adversity. Mitochondrion, 57, 213–221.

