## Supplemental Material for "Glutamatergic and Redox Abnormalities in Major Depressive Disorder: Links to Childhood Maltreatment and Perceived Stress"

***Supplementary Material***

**Supplementary Methods**

- Detailed inclusion and exclusion criteria
- Participant- and variable-specific exclusions
- Internal consistency of clinical measures

**Supplementary Methods**

**Detailed inclusion and exclusion criteria**

Inclusion criteria for all participants:

- Female sex assigned at birth

- Ages from 20 to 32

- Right- or left-handed^*^

- Capable of providing written informed consent

- Fluency in written and spoken English

- Absence of current dopaminergic medication use in the past 3 months, including norepinephrine and dopamine reuptake inhibitors (NDRIs) such as bupropion and psychostimulants such as Adderall and methylphenidate

- No evidence of significant inconsistencies in self-reports

Exclusion criteria for all participants:

- Lifetime history of psychosis or mania

- Past/current DSM-5 diagnosis of: OCD, schizophrenia, schizoaffective disorder, delusional disorder, psychotic disorders NOS, bipolar disorder, patients with mood congruent or mood incongruent psychotic features, autism or any other pervasive developmental disorder, organic mental disorder

- Current diagnosis of PTSD; however, in patient groups a history of PTSD will be allowed if the symptoms have been subthreshold for the past month

- Specific phobia, panic disorder, social anxiety disorder, and generalized anxiety disorder allowed in patient groups only if judged to be currently mild and never the principal diagnosis when co-occurring with current or past MDD

- No current mild, moderate, or severe alcohol or substance use disorder. Past mild, moderate, or severe alcohol or substance use disorder is allowed as long as no criteria for any level of abuse have been met in the past 3 months, i.e., the participant is in early or sustained remission (except for any history of cocaine or stimulant abuse, which will lead to automatic exclusion)

- Recent history of anorexia (past 1 year); or bulimia (past 3 months)

- Lifetime history of neurological impairment, significant head injury, or concussion

- MRI contraindications

- Pregnant women or women who gave birth in the last 12 weeks

- Patient is clinically unstable, in the judgment of the clinician

For individuals within the remitted MDD group, the following additional criteria were required:

- At least one major depressive episode (MDE) within the past five years

- MDD in remission for at least two months

- No more than two symptoms of depression reported to more than a mild degree (SCID rating of 2) in the eight weeks prior to testing. Current depressed mood and anhedonia symptoms (SCID criteria #1 and #2) must be rated 1

- Does not meet criteria for Current Persistent Depressive Disorder

For healthy controls, participants had to be free of any lifetime medical, neurological, and psychiatric illness (including alcohol/substance abuse) and had no first-degree relatives with mood disorders or psychosis.

^*^Eligibility criteria permitted both right- and left-handed participants; however, only one participant in the final sample was left-handed.

**Participant- and variable-specific exclusions**

**Self-report reliability.** MACE severity and multiplicity scores were excluded for three participants in the MDD/CSA group whose self-reported maltreatment histories were discordant with clinical interview. Two disclosed CSA during clinical interview but did not endorse the corresponding exposure on self-report measures; the third endorsed CSA on the MACE but not on a second self-report instrument and, when queried during the clinical interview, reported not having attended fully to the questionnaires. In each case, the clinician confirmed CSA history, diagnostic group assignment followed the clinical determination, and self-report maltreatment scores were treated as unreliable.

**Statistical outliers.** Outliers were defined as observations exceeding three standard deviations from the full-sample mean. The following outliers were excluded: glutamate (*n* = 1, HC), NAD⁺/NADH (*n* = 1, MDD/CSA), and MACE severity (*n* = 1, rMDD/CSA). The participant excluded from MACE severity analyses was retained in the MACE multiplicity analyses because the raw multiplicity score fell within the three-standard-deviation threshold calculated for the CSA-exposed subsample, whereas her severity score exceeded that threshold as well as the full-sample threshold.

**Missing data.** Glutamate, glutamine, and Glx were unavailable for two participants in the rMDD/CSA group because of ¹H-MRS acquisition failure; their ³¹P-MRS measurements were retained. NAD⁺/NADH data were unavailable for three additional participants. Missing 6-month perceived-stress assessments reflected incomplete follow-up rather than outlier exclusion. Resulting sample sizes were: glutamate n = 68; glutamine and Glx n = 69; NAD⁺/NADH n = 67; MACE severity n = 26 and multiplicity n = 27 (CSA-exposed subsample); BDI-II n = 28 (currently depressed subsample); perceived stress n = 67 and 61 at baseline and 6-month follow-up.

**Internal consistency of clinical measures**

Cronbach’s α for each clinical measure in the analyzed sample is presented by group in Table S1.

**Table S1**

*Cronbach’s α of Clinical Measures by Group*

| Measure | HC | MDD/no CSA | MDD/CSA | rMDD/CSA | Total |
| --- | --- | --- | --- | --- | --- |
| PSS-14, baseline | .84 (25) | .64 (12) | .81 (12) | .81 (15) | .95 (64) |
| PSS-14, 6 months | .78 (26) | .90 (12) | .71 (10) | .81 (13) | .94 (61) |
| BDI-II | — | .69 (14) | .86 (13) | — | .83 (27) |

*Note.* Values are Cronbach’s α (*n* with complete item data). The BDI-II was analyzed among participants with current MDD only. CSA = childhood sexual abuse; HC = healthy controls; MDD = major depressive disorder; rMDD = remitted major depressive disorder; PSS-14 = 14-item Perceived Stress Scale; BDI-II = Beck Depression Inventory-II.
